# Systems medicine links microbial inflammatory response with glycoprotein-associated mortality risk

**DOI:** 10.1101/018655

**Authors:** Scott C Ritchie, Peter Würtz, Artika P Nath, Gad Abraham, Aki S Havulinna, Antti J Kangas, Pasi Soininen, Kristiina Aalto, Ilkka Seppälä, Emma Raitoharju, Marko Salmi, Mikael Maksimow, Satu Männistö, Mika Kähönen, Markus Juonala, Terho Lehtimäki, Sirpa Jalkanen, Markus Perola, Olli Raitakari, Veikko Salomaa, Mika Ala-Korpela, Johannes Kettunen, Michael Inouye

**Affiliations:** Centre for Systems Genomics, School of Biological Sciences, The University of Melbourne, Parkville 3010, Victoria, Australia; Department of Pathology, The University of Melbourne, Parkville 3010, Victoria, Australia; Computational Medicine, Institute of Health Sciences, University of Oulu, Oulu, Finland; National Institute for Health and Welfare, Finland; Institute for Molecular Medicine Finland, University of Helsinki, Helsinki, Finland; NMR Metabolomics Laboratory, School of Pharmacy, University of Eastern Finland, Kuopio, Finland; MediCity Research Laboratory, Department of Medical Microbiology and Immunology, University of Turku, Turku, Finland; Department of Clinical Chemistry, Fimlab Laboratories, University of Tampere, School of Medicine, Tampere, Finland; Department of Clinical Physiology, University of Tampere and Tampere University Hospital, Tampere; Department of Medicine, University of Turku and Division of Medicine, Turku University Hospital, Turku, Finland; Murdoch Childrens Research Institute, Parkville, Victoria, Australia; Department of Clinical Physiology and Nuclear Medicine, Turku University Hospital, Turku, Finland; Research Centre of Applied and Preventive Cardiovascular Medicine, University of Turku, Turku, Finland; Oulu University Hospital, Oulu, Finland; Computational Medicine, School of Social and Community Medicine, University of Bristol, Bristol, United Kingdom; Medical Research Council Integrative Epidemiology Unit at the University of Bristol, Bristol, United Kingdom

## Abstract

Integrative analyses of high-throughput omics data have elucidated the aetiology and pathogenesis for complex traits and diseases^1–4^, and the linking of omics information to electronic health records promises new insights into human health and disease. Recent nuclear magnetic resonance (NMR) spectroscopy biomarker profiling has implicated glycoprotein acetyls (GlycA) as a biomarker for cardiovascular risk^5^ and all-cause mortality^6^. To elucidate biological processes contributing to GlycA-associated mortality risk, we leveraged human omics data from three population-based cohorts together with nation-wide Finnish hospital and mortality records. Elevated GlycA was associated with myriad infection-related inflammatory processes. Within individuals, elevated GlycA levels were stable over long time periods, up to a decade, and chronically elevated GlycA was also associated with modest elevation of numerous cytokines. Individuals with elevated GlycA also showed increased expression of a transcriptional sub-network, the Neutrophil Degranulation Module (NDM), suggesting an increased activity of microbe-driven immune response. Subsequent analysis of nation-wide hospitalisation and death records was consistent with a microbial basis for GlycA-associated mortality, with each standard deviation increase in GlycA raising an individual’s future risk of hospitalization and death from non-localized infection by 40% and 136%, respectively. These results show that, beyond its established role in acute-phase response^7–9^, elevated GlycA is more broadly a biomarker for low-grade chronic inflammation and increased neutrophil activity. Further, increased risk of susceptibility to severe microbial-infection events in healthy individuals suggests this inflammation is a contributor to mortality risk. Taken together, this study demonstrates the power of an integrative approach that combines omics data and health records to delineate the biological processes underlying a newly discovered biomarker, providing a model strategy for future systems medicine studies.

## Main text

The NMR signal for GlycA has long been associated with acute-phase inflammatory response^7–9^, but has recently been the subject of intense research due to its predictive capacity for incident cardiovascular disease risk^5^ and all-cause mortality^6^. Here, we aim to elucidate the molecular pathways underlying the elevated disease and mortality risk associated with GlycA (study workflow given in **Figure S1**). It is well-known that GlycA is a complex heterogeneous NMR signal which reflects the abundance of *N*-acetyl groups located on glycoproteins in circulation^7,10,11^. Many abundant proteins in circulation are glycoproteins and the majority are acute-phase proteins: rising or falling in response to myriad perturbations, including inflammation, infection, drug usage, and tissue injury^12^. To fine-map the GlycA NMR signal, we assayed four abundant acute-phase glycoproteins (alpha-1-acid glycoprotein, haptoglobin, alpha-1-antitrypsin, and transferrin) in serum samples from the population-based Dietary, Lifestyle, and Genetic determinants of Obesity and Metabolic syndrome study (DILGOM)^13^ (**Methods**). Consistent with previous studies^7,10,11^, we found that all four glycoproteins were significantly associated with GlycA (**Table S1**). Variation in GlycA was best explained by levels of alpha-1-acid glycoprotein, followed by haptoglobin, transferrin, and alpha-1-antitrypsin. The rank of the associations was in accordance with previous estimates of *N-*acetyl side-chain proportions in each protein^10^, as well as previous observations that alpha-1-acid glycoprotein is the most abundant glycoprotein in circulation^6,10,11^.

The association of GlycA with concentrations of multiple circulating glycoproteins is consistent with the recognised role of GlycA as a summary measure of acute-phase glycoproteins as well as the suggestion that, in the context of mortality risk, it is a marker for systemic inflammation^5,6^. Yet, the current clinical standard for assessing acute-phase and chronic systemic inflammation, C-reactive protein (CRP)^14^, does not markedly change the prediction models for cardiovascular disease events^5^ or all-cause mortality^6^, suggesting that GlycA and CRP capture partly different aspects of inflammatory response. We investigated the relationship between GlycA and inflammatory response in the longitudinal population-based Cardiovascular Risk in Young Finns Study (YFS)^15^ (**Methods**). First, consistent with its sensitivity to acute-phase response^7–9^, we observed a significant elevation of GlycA in individuals who reported an infection with fever in the two weeks prior to sample collection (mean increase of 0.41 standard deviations; **Figure S2**). Next, we utilized multiplex cytokine panels on serum samples from YFS participants without recent self-reported infection to investigate the relationship between GlycA and circulating immune signalling molecules (**Methods**). GlycA is known to interact with IL-6^10,12^, however we observed a large number of robust associations (Bonferroni-adjusted P < 0.001): 32 of 36 tested cytokines were significantly associated with GlycA (**Figure 1**, **Table S2**). The levels of 29 cytokines, with both anti- and pro-inflammatory roles, were positively associated with GlycA (**Figure 1**). In contrast, only Cutaneous T-cell-attracting chemokine (CTACK), showed a prominent negative association with GlycA (**Figure 1**). Although there were numerous significant associations, no single cytokine showed a correlation with GlycA that alone could explain its association with mortality or cardiovascular disease risk. To investigate the dynamics of GlycA levels outside of acute-phase, we utilized the GlycA time-series available for YFS participants (2001, 2007, and 2011), finding that within individuals there was a strong auto-correlation of GlycA levels across all three time points (mean Pearson *r* = 0.43; **Table S3**). These results provide evidence for the association of GlycA with systemic inflammation, while further showing its chronicity and broad summarization of myriad pro- and anti-inflammatory cytokines in circulation.

**Figure 1:**
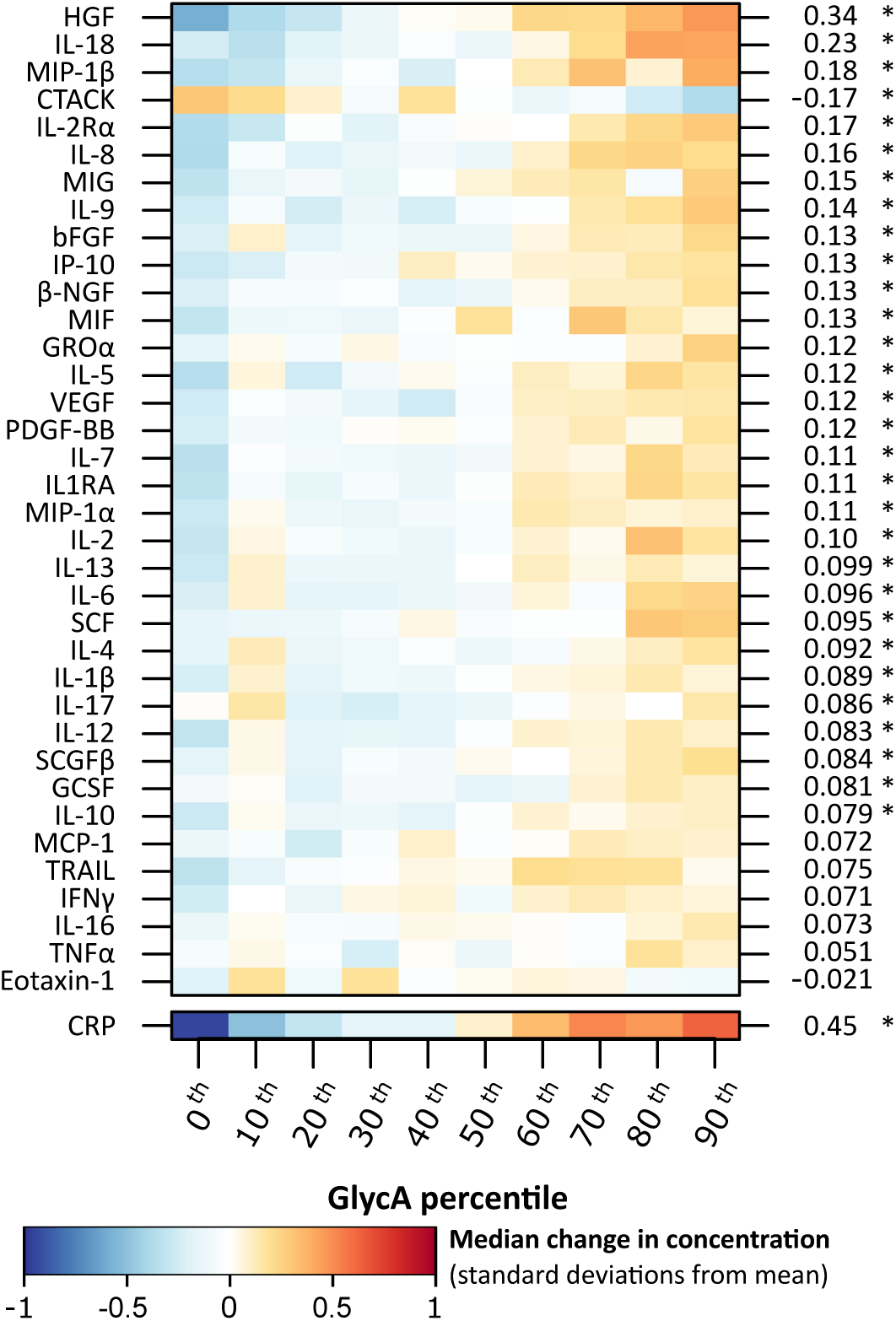
GlycA summarises low-grade inflammation cytokine activity. Median difference in concentration (standard deviations) relative to the population mean for 36 assayed cytokines (**Methods**) and C-reactive protein (CRP) for ten GlycA-risk categories in the 2007 collection of the YFS cohort. The right axis denotes the change in SD-units of GlycA per standard deviation (SD) increase of each cytokine or CRP adjusting for age and sex in the YFS cohort (**Table S2**). *: The association is significant at P < 0.001 (Bonferroni corrected for the number of cytokines).

To gain insight into the cellular processes associated with elevated circulating GlycA, we utilized whole blood transcriptome profiling in the DILGOM and YFS cohorts. Weighted transcriptional networks^16^ were constructed in the DILGOM cohort and then tested for association with GlycA levels, with significantly associated subnetworks tested for replication in the YFS cohort using permutation of network topology features, such as edge density and connectivity^17^ (**Methods**). We reconstructed a total of 40 transcriptional sub-networks (or modules), of which three were significantly associated with GlycA after adjustment for multiple testing (adjusted P < 0.001; **Table S4**). The models were corrected for age, sex and additionally with triglyceride levels, as in the previous study for all-cause mortality we observed a multivariable effect when serum triglyceride measures as captured by VLDL diameter and GlycA were in the same models^6^; thus the multivariable effect was here used to focus on GlycA-specific variation and its contribution to inflammatory mortality risk. The GlycA association and network topologies for two (Modules A and B) of the three modules replicated in the YFS cohort (**Table S5**). Although Module B replicated (**Figure S3**), its inferred biological function was incoherent; no significant KEGG pathways were identified and the majority of enriched GO terms (28 / 39) were attributable to a non-central gene (**Table S6**). In contrast, Module A represented stronger GlycA association, replication, and clear biological function (**Figure 2**, **Tables S7-8**).

**Figure 2:**
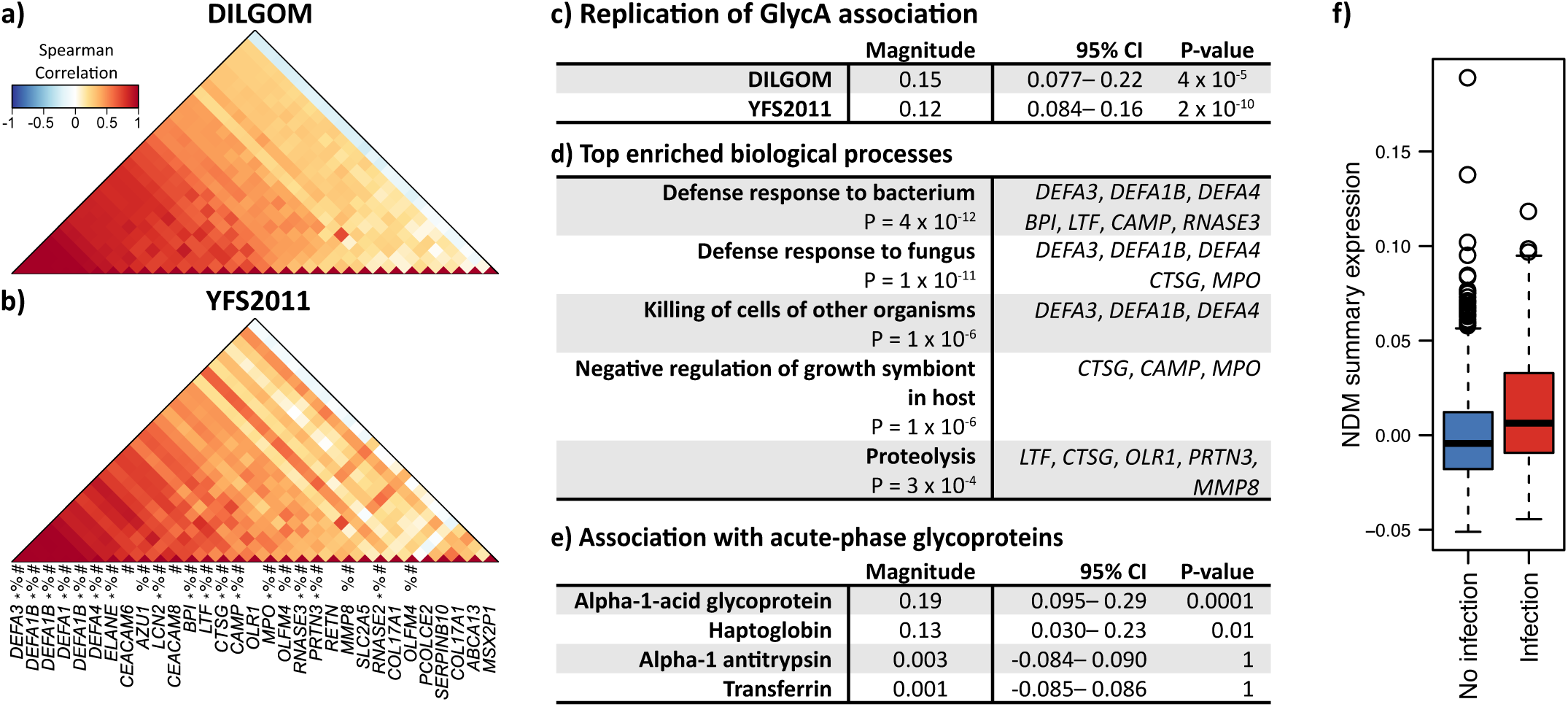
The Neutrophil Degranulation Module (NDM). Probe coexpression (Spearman correlation) in the DILGOM cohort (**a**) and replication in the YFS2011 cohort (**b**). See **Table S9** for module gene details. Symbols above gene symbol indicate expression in neutrophils (#), whether its product is located in neutrophil granules (%), and whether its product has antimicrobial functionality (*) (**Table S10**). **c)** Replication of GlycA association in both cohorts. Associations were assessed using a linear regression of GlycA on the module summary expression adjusting for age, sex, and triglycerides (**Methods**). Magnitude corresponds to change in SD-units of GlycA per SD increase of NDM summary expression. CI: confidence interval. **d)** The top enriched Gene Ontology Biological Process terms. A full list of significant enrichments can be found in **Table S8**. **e)** Association of NDM module summary expression with alpha-1 acid glycoprotein, haptoglobin, alpha-1 antitrypsin, and transferrin in the DILGOM cohort in a multivariable linear regression (**Methods**). Magnitude corresponds to change in SD-units of GlycA per SD increase of each respective acute-phase glycoprotein. **f)** Association with self-reported febrile infection (N = 72 with infection, N = 1,550 without infection) in the two weeks prior to blood sample collection in the YFS2011 cohort. The mean difference in NDM summary expression was 0.61 (95% confidence interval: 0.29–0.93, P = 3 x 10^-4^ from a t-test; **Methods**). GlycA and triglyceride levels were log transformed. All continuous measurements were standardised.

Comprising 27 genes (**Table S9**), Module A showed a prominent neutrophil transcriptional signature (19 genes, or 70% expressed in neutrophils) as well as a functional role in neutrophil degranulation, with 17 genes (63%) coding for products located in neutrophil granules, 14 (52%) of which serve antimicrobial functions (**Figure 2**; **Table S10**)^18–21^. The enriched KEGG pathways and GO processes largely implicated innate immune response to microbes (**Figure 2d, Tables S7-8**). Supporting this, Module A summary expression (**Methods**) was elevated by 0.61 standard deviations (P = 3 x 10^-4^) in individuals reporting febrile infection within 2 weeks prior to blood sampling (**Figure 2f**) and there were robust positive associations between Module A summary expression profile and concentrations of alpha-1-acid glycoprotein and haptoglobin (**Figure 2e**). Although alpha-1-acid glycoprotein and haptoglobin are primarily synthesised by hepatocytes in the liver, they have also been found to be synthesised in, and secreted from, neutrophil granules^22^. To test for genetic control of Module A expression, we performed a transcriptional module QTL scan (**Methods**). We identified three genome-wide significant genetic loci (**Table S11a**), the top signal of which was a SNP intronic to *LRRC8A* (rs13297295). This module QTL was a strong driver of *LCN2*, 750kb upstream, a gene of high centrality to the module (P = 4 x 10^-10^; **Table S11b**) and whose Toll-like receptor induced protein product has pivotal bacteriostatic properties^18^, particularly through sequestration of bacterial siderophores which scavenge iron^23^. Given these results overall, we subsequently refer to this module as the Neutrophil Degranulation Module (NDM).

Neutrophils are important first responders in innate immune response to invading microbes. Once recruited to the site of infection, neutrophils employ a number of mechanisms to degrade and destroy microbes, one of which is degranulation: the exocytosis of antimicrobial peptides stored within their granules^22^. Importantly, neutrophil antimicrobial peptides are also cytotoxic; they damage and degrade tissue at the site of their release. The production and influx of neutrophils must therefore be carefully regulated^22^. To investigate the relationship between NDM expression and neutrophil abundance, we utilized white blood cell count data from whole blood in YFS. We observed a strong positive association between leukocyte abundance and elevated NDM expression (P = 2 x 10^-17^; **Table S12**). Neutrophils account for 40– 80% of leukocytes in circulation, suggesting that elevated NDM expression is to some extent associated with increased circulating neutrophil abundance. In models considering both NDM expression and leukocyte abundance, both displayed strong independent associations with GlycA levels (P = 2 x 10^-6^ and P = 2 x 10^-15^ respectively; **Table S13**), indicating that elevated GlycA is associated with upregulation of neutrophil activity through both increased circulating neutrophil populations as well as increased expression of neutrophil-derived cytotoxic peptides.

To investigate whether chronic elevation of microbial-mediated inflammatory response may result in an increased risk for infection-associated morbidity and mortality, we analysed electronic hospital records over a 13.8-year follow-up period for 7,599 individuals with GlycA profiles from the population-based FINRISK97 cohort (**Methods**). We observed a markedly increased risk of both hospitalization and death from severe microbial infection for individuals with elevated GlycA (**Figure 3**). Strikingly, individuals with GlycA above the population median were at more than 5-fold increased risk of death from non-localized infections (**Figure 3a**). Each increment of one standard deviation in GlycA concentration conferred more than two-fold higher risk of death from non-localized infections, largely attributable to death from septicaemia, and a 38% increase risk of death from pneumonia (**Figure 3b**). GlycA was also more broadly predictive for hospitalization from infection, including non-localized bacterial and fungal infections, respiratory infections, and localized infections to the skin, bones, joints, intestinal, and urinary system (**Figure 3b**). Compared to CRP, the hazard ratios for GlycA were consistently higher (**Figure S4**) and only weakly attenuated when adjusting the prediction models for CRP (**Figure S5**).

**Figure 3:**
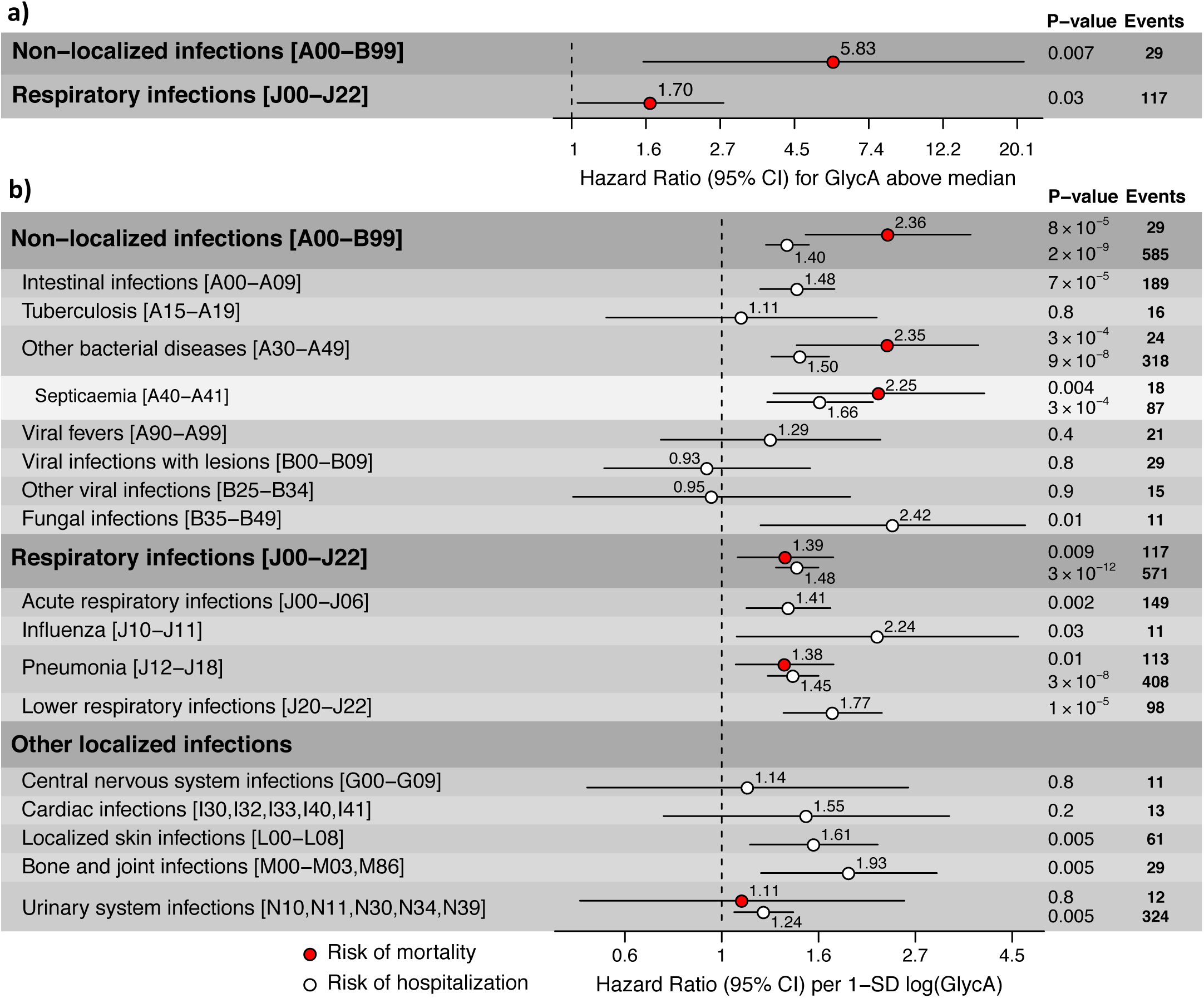
GlycA predicts hospitalisation and death from infectious diseases. Hazard ratios for fatal events from broad infection-related categories for (**a**) individuals with GlycA concentrations above the population median, and (**b**). risk of fatal events (red) or hospitalization (white) conferred per standard deviation increment of log transformed GlycA for infection-related diagnoses with more than 10 events in the FINRISK97 cohort. 7,599 apparently healthy individuals from the general population were prospectively observed over a 13.8-year follow-up period. Cox models were adjusted for age, sex, triglycerides and incidence of the same diagnosis in the 10-years prior to sample collection. Numbers in square brackets indicate the ICD-10 code blocks for the corresponding diagnosis. See **Figure S6** for an extended breakdown into individual ICD-10 diagnosis codes.

The increased risk of severe infection is consistent with overactivity of anti-microbial immune response and reflective of the elevation in cytokine levels. While no one cytokine was particularly associated with GlycA, it was notable that a diverse range of pro- and anti-inflammatory cytokines showed a consistent positive association, indicating that GlycA is able to capture broad changes to circulating cytokine patterns. Overactivity of immune response from severe infection events, such as sepsis and influenza, is known to cause tissue damage and organ dysfunction through hypercytokinemia^24^. Our transcriptomic analyses indicate that GlycA-related immune overactivation may also occur through the NDM and increased neutrophil activity. Taken together, these results suggest that anti-inflammatory treatments may have some effect on ameliorating GlycA-associated mortality risk. In particular, consistent with our observations that each standard deviation increase in GlycA was associated with 38% higher risk of death from pneumonia in an apparently healthy population (**Figure 3**), it has been suggested previously that corticosteroids can in some cases reduce risk of treatment failure for hospitalized patients with severe pneumonia and excessive inflammatory response^25^.

This study sheds light on biological processes underlying GlycA-associated mortality risk, elucidating low-grade, infection-related inflammation as a likely driver of increased mortality risk from increased susceptibility to severe infection later in life. Our study represents a potentially useful strategy for future integration of multi-omics data with electronic health records for uncovering the biological basis for novel disease biomarkers.

## Methods

### Cohort and sample descriptions

All study participants provided written informed consent. Protocols were designed and performed according to the principles of the Helsinki Declaration, and data protection, anonymity and confidentiality have been assured.

The Cardiovascular Risk in Young Finns Study (YFS) is an ongoing population-based prospective cohort study across five major population centres in Finland (Helsinki, Kuopio, Turku, Oulu, and Tampere). The baseline study began in 1980, recruiting 4,320 children and adolescents randomly from the population in six age groups: 3, 6, 9, 12, 15, and 18 years of age. A total of 3,596 participated in the original study. For this study the three most recent follow-up studies were utilized. A total of 2,283, 2,202, and 2,063 subjects participated in the clinical examinations in 2001, 2007, and 2011 respectively. Drop-out rates have been dynamic: some individuals have dropped out only to re-participate in later collections, however participants at each time point were representative of the baseline cohort. Ethics was approved by the Joint Commission on Ethics of the Turku University and the Turku University Central Hospital ^15^. Not all collection years were available for molecular profiling, thus some association analyses within years were not possible. Details of the molecular profiling are given below.

The Dietary, Lifestyle, and Genetic determinants of Obesity and Metabolic syndrome (DILGOM) study is a cross-sectional population survey conducted in 2007 as an extension of the FINRISK 2007 study. In the greater Helsinki region, the study randomly recruited 631 unrelated Finnish individuals aged 25–74. Ethics was approved by the Coordinating Ethical Committee of the Helsinki and Uusimaa Hospital District^13^.

The FINRISK study is an ongoing cross-sectional population survey of Finland recruiting a random sample of 6,000–8,800 individuals every five years since 1972. This study utilized the 1997 collection (FINRISK97), which recruited 8,444 individuals aged 25–74 from five major regional and metropolitan areas of Finland: the provinces of North Karelia, Northern Savo, and Northern Ostrobothnia and Kainuu; the Turku and Loimaa region of south-western Finland; and the Helsinki and Vantaa metropolitan area. Ethics was approved by the ethical committee of the National Public Health Institute, Finland ^26^.

Blood sample collection protocols were similar across the three cohorts analysed in this study and have been described previously^13,15,26^. Briefly, a blood sample was taken and extensive clinical characteristics and lifestyle information collected from subjects participating in each cohort. Blood samples were collected after an overnight fast from the DILGOM and YFS cohorts. Non-fasting blood samples were collected for the FINRISK97 cohort. Serum samples were subsequently aliquoted and stored at -70°C.

### Molecular profiling

GlycA, CRP, and total triglycerides were quantified from serum in the DILGOM (N = 583), YFS2001 (N = 2,245), YFS2007 (N = 2,159), YFS2011 (N = 2,046), and FINRISK97 (N = 7,599) studies. GlycA and total triglycerides were quantified using a high-throughput proton NMR metabolomics platform. Experimental protocols including sample preparation and spectroscopy were as previously described^27^. High-sensitivity serum CRP was measured by an automated analyser using a latex turbidimetric immunoassay kit.

Concentrations for alpha-1-acid glycoprotein, haptoglobin, alpha-1 antitrypsin and transferrin were quantified from serum using modular analysers and Roche Tina-quant assays using the immunoturbidimetric method for 630 individuals in the DILGOM cohort. The intra-individual coefficient of variation was < 3% for all four assays.

A panel of 48 cytokines was quantified from serum samples for 2,200 individuals in YFS2007 on Bioplex multiplex cytokine assays. The Bio-Rad analyser program was used to fit concentrations to standard curves for each cytokine on each 96-well plate using a five-parameter logistic regression. Due to the logistic, non-linear standard curves, the upper and lower detection limits were calculated plate-wise, so that they corresponded with “asymptotic” concentrations representing fluorescent intensity 2% above lower and 2% below upper asymptote of the calibration curve. Cytokines were excluded from further analysis if more than 10% of their observations were missing or corresponded to asymptotic concentrations (12 cytokines excluded).

Genome-wide expression profiling was carried out for the DILGOM (N = 518) and YFS2011 (N = 1,650). RNA was quantified from peripheral blood as previously described^13,28^ and hybridized to the Illumina HT12 version 3 (DILGOM) and version 4 (YFS2011) BeadChip arrays. Expression arrays were processed as previously described for DILGOM^13^ and YFS2011^28^. Briefly, DILGOM arrays were quantile normalised at the strip level, and technical replicates were combined by a weighted bead-count average or removed if their concordance was low^13^. YFS2011 arrays were processed using nonparametric background correction, quantile normalised, and log_2_ transformed^28^.

Genome-wide genotyping was carried out for the DILGOM (N = 518) and for the YFS (N = 2,443). DNA was extracted from whole blood samples as previously described^13^, and genotyped on an Illumina 610-Quad SNP array (DILGOM)^13^ and a custom 670K Illumina BeadChip array sharing 562,643 SNPs in common with the Human610 BeadChip array (YFS)^29^. Genotype calling and quality control procedures were as previously described^13,29^. Missing genotypes were imputed to the 1000 genomes reference panel and imputed SNPs were filtered based on low call-rate (< 0.90 for DILGOM, < 0.95 for YFS), low-information score (< 0.4), minor allele frequency < 1%, deviation from Hardy-Weinberg equilibrium (P < 5.00 x 10^-6^). 7,263,701 and 6,721,082 SNPs passed quality control in the DILGOM and YFS cohorts, respectively.

Blood cell count analysis was carried out for 2,027 individuals from the YFS2011 cohort. Venous blood was anticoagulated with EDTA and measured by flow cytometric particle counting (Sysmex Corporation, Kobe, Japan) with reagents provided by the manufacturer (Cellpack and Sulfolyser).

### Hospital discharge and causes-of-death records

Nationwide diagnosis data was obtained from the Finnish National Hospital Discharge Register and the National Causes-of-Death Register for the FINRISK97 participants with GlycA measured (N = 7,599)^30^. Diagnoses data was obtained for a 13.8-year follow-up period, and for 10 years prior to sample collection. Diagnoses from 1997 to 2011 were encoded according to the International Classification of Diseases (ICD) 10^th^ Revision (ICD-10). Diagnoses occurring from 1987 to 1996 were encoded in ICD 9th Revision (ICD-9) format and converted to ICD-10 format by the scheme provided by the US Center for Disease Control Diagnosis Code Set General Equivalence Mappings (ftp://ftp.cdc.gov/pub/Health_Statistics/NCHS/Publications/ICD10CM/2011/). Diagnosis conversions were further verified according to the mapping scheme provided by the New Zealand Ministry of Health; National Data Policy Group (http://www.health.govt.nz/nz-health-statistics/data-references/mapping-tools/mapping-between-icd-10-and-icd-9). Manual curation of the ICD-9 to ICD-10 conversion was conducted for all diagnoses with mismatch to the degree of 3 digits between the two conversion protocols.

### Network analysis

Weighted gene coexpression network analysis (WGCNA)^16^ (version 1.42) was used to identify modules of coexpressed probes within the whole blood gene expression of the DILGOM cohort. To ensure that differences of age or sex on gene expression did not influence coexpression inference and module detection, probe intensities were adjusted by taking the residuals from a multivariable linear regression of each probe on age and sex prior to network construction. We used the recommended parameters for WGCNA, except for the following: Spearman’s rank correlation coefficient was used as the measure of pairwise-probe coexpression and a minimum module size of 10 was specified. The soft-threshold power selected by the automated scale-free topology criterion procedure was 5. Summary expression profiles for each module were calculated as the first eigenvector from a principal component analysis on the module’s gene expression.

Replication of module topology (**Table S5a**) in the YFS2011 was assessed through a permutation procedure using previously described module preservation statistics^17^. Briefly, coexpression and adjacency matrixes were calculated for the YFS2011 as described for DILGOM. A soft-threshold power of 4 was selected by automatic selection procedure in the YFS2011 cohort. The module preservation statistics were then calculated on 1,000,000 random permutations of module assignments in YFS2011, yielding a permuted P-value. Under this framework, the minimum P-value obtainable was 1 x 10^-6^. A module’s topology was considered replicated if all module preservation statistics were significant at P < 0.05. The mean intramodular probe connectivity (**Table S9**) for Module A, the neutrophil degranulation module (NDM) was calculated as the sum of the adjacencies to other module probes averaged across both the DILGOM and YFS2011 cohorts.

Over-representation analysis of Gene Ontology (GO) Biological Process terms (**Figure 2d, Table S6**, **S8**) and KEGG Pathway terms (**Table S7**) for each replicable GlycA-associated module was performed using the web-tool, GeneCoDis^31^. An annotation was considered significant if its FDR corrected hypergeometric p-value was < 0.05.

### Statistical analysis

A natural log transform was applied to GlycA, each glycoprotein, and triglycerides. A rank based inverse normal transform was applied to each cytokine. After transformations, all continuous measurements (GlycA, glycoproteins, CRP, cytokines, triglycerides, and network module summary expression profiles) were standardised to SD units (standard deviations) throughout.

A multivariable linear regression model adjusting for age and sex was used to assess associations between GlycA and the four glycoproteins assayed in the DILGOM cohort (**Table S1**).

A t-test was used to assess the association between GlycA and self-reported febrile infection in the two weeks prior to blood sampling (**Figure S2**). Measurements for GlycA and self-reported infection were available for all three adulthood surveys of the YFS cohort (2001, 2007, and 2011). Reported differences indicate the mean elevation of GlycA in SD-units for those reporting febrile infection compared to those reporting no febrile infection.

A linear regression model was used to assess association of GlycA with each cytokine and CRP for participants from the YFS2007 cohort reporting no febrile infection within the two weeks prior to blood sample collection (**Figure 1**, **Table S2**). Models were adjusted for age and sex. Association magnitudes denote the difference in SD-units of GlycA per SD increase of each cytokine or CRP. Associations between GlycA and each cytokine were considered significant at a Bonferroni adjusted threshold of P < 0.001.

A linear regression model was used to assess associations between GlycA within participants longitudinally using time-series available for the YFS cohort (2001, 2007, and 2011; **Table S3**). Models were adjusted for age and sex. Association magnitudes denote the difference in SD-units of GlycA per SD increase of GlycA at a previous survey.

Associations between GlycA and the 40 modules reconstructed in the DILGOM cohort were assessed through linear regression of GlycA on each module’s summary expression profile (Principal Component 1 from data scaled to SD-units) adjusting for age, sex, and triglyceride levels; as in the previous study for all-cause mortality we observed a multivariable effect when serum triglyceride measures as captured by VLDL diameter and GlycA were in the same models^6^ (**Table S4**). Association magnitudes correspond to change in standard deviation of GlycA per standard deviation increase in each module’s summary expression profile. An association between GlycA and a module was considered significant at a Bonferroni corrected threshold of P < 0.001 (adjusting for the number inferred modules). The variance explained by each module’s summary expression profile was also reported (**Table S4**). Associations with GlycA were further tested in the YFS2011 cohort (as described above) for modules that were significantly associated with GlycA in the DILGOM cohort (Modules A (NDM), B, and C; **Table S5b**).

A multivariable linear regression was used to assess associations between NDM summary expression and the four glycoproteins assayed in the DILGOM cohort (**Figure 2e**). Glycoproteins were only assayed for the DILGOM cohort. Association magnitudes denote the difference in SD-units of NDM summary expression per SD increase of each protein.

A t-test was used to assess the association between the summary expression (Principal Component 1) of the NDM and self-reported febrile infection in the two weeks prior to sampling in the YFS2011 cohort (**Figure 2f**). Self-reported infection status was only available for the YFS cohort. NDM summary expression was standardised in the model. Reported differences indicate mean elevation of NDM summary expression in SD-units for participants reporting febrile infection compared to those reporting no febrile infection.

Quantitative trait loci (QTLs) for the neutrophil degranulation module (NDM) were identified through a genome-wide scan performed for the summary expression profile of the NDM with meta-analysis of the DILGOM and YFS2011 cohorts using a fixed-effects model (**Table S11a**). Genome-wide association testing and meta-analysis were performed using plink (v1.0.7) software. Individual associations were tested using a linear model of SNP dosage on NDM summary expression. A SNP was considered a QTL for the NDM if the P-value from the meta-analysis was less than a genome-wide significance threshold of P < 5 x 10^-8^. The models for the genome-wide association study in the both cohorts were adjusted for the first two principal components of the genotype data to control for population structure. The module QTL rs13297295 was further tested for association with the *cis*-module gene *LCN2* (750kB upstream) in the YFS2011 cohort (**Table S11b**).

Linear regression models were used to test the association between leukocyte abundance and the NDM summary expression profile (**Table S12**). White blood cell counts were available only the YFS2011 cohort. Association magnitudes corresponded to difference in SD-units of NDM summary expression per SD increase of leukocyte abundance. A multivariable linear regression model was further used to test if leukocyte abundance and the NDM summary expression profile were independently associated with GlycA (**Table S13**). Regression models were fitted with age, sex, and triglycerides as covariates. Association magnitudes corresponded to change in SD-units of GlycA per standard deviation increase of leukocyte abundance or NDM summary expression.

GlycA was tested for association with incidence of infectious disease leading to hospitalization or death in the FINRISK97 cohort over a 13.8-year follow-up period resulting in prospective observation during 99,671 person-years. ICD-block structures, and specific diagnoses within the ICD-10 categories A00–B99 (non-local infections), J00–J22 (respiratory infections), and other localized infections (G00-G09, I30, I32, I33, I40, I41, L00–L08, M00–M03, M86, N10, N11, N30, N34, and N39) with more than 10 incident events were analysed (**Figure 3, Figure S6**). Hazard ratios of GlycA for hospitalization or death were assessed by Cox proportional-hazard models using age as the time-scale and adjusted for sex, triglycerides and prior occurrence of the pertinent diagnosis category within the 10 years prior to blood sample collection. The models were further tested by additionally adjusting for CRP (**Figure S5**), and by replacing GlycA for CRP (**Figure S4**).

Unless otherwise stated, all analyses were performed using the statistical computing software, R version 3.1.1, (http://www.r-project.org/).

## Acknowledgements

This study was supported by funding from National Health and Medical Research Council of Australia (NHMRC) grant APP1062227. MI was supported by an NHMRC and Australian Heart Foundation Career Development Fellowship (no. 1061435). SR was supported by an Australian Postgraduate Award and a PhD student top-up award from Victorian Life Sciences Computation Initiative (VLSCI).

This study was further supported by the Strategic Research Funding from the University of Oulu, Finland, the Sigrid Juselius Foundation, the Academy of Finland (grant number 141136), the Yrjö Jahnsson Foundation, the Emil Aaltonen Foundation, and the Finnish Diabetes Research Foundation. The Cardiovascular Risk in Young Finns Study is supported by the Finnish Foundation for Cardiovascular Research, Oulu, Helsinki, Kuopio, Tampere, and Turku University Central Hospital Medical Funds, the Paavo Nurmi Foundation, the Juho Vainio Foundation, the Finnish Cultural Foundation, the Finnish Funding Agency for Technology and Innovation. MJ was supported by the Paulo Foundation, Maud Kuistila Foundation, and Finnish Medical Foundation. The DILGOM study and the National FINRISK study are supported by the Academy of Finland (grant numbers 139 635).

The quantitative serum NMR metabolomics platform and its development has been supported by the Academy of Finland, TEKES - the Finnish Funding Agency for Technology and Innovation, the Sigrid Juselius Foundation, the Novo Nordisk Foundation, the Finnish Diabetes Research Foundation, the Paavo Nurmi Foundation, and the strategic and infrastructural research funding from the University of Oulu, Finland, as well as by the British Heart Foundation, the Wellcome Trust, and the Medical Research Council, UK.

